# Music training improves the ability to understand speech-in-noise in older adults

**DOI:** 10.1101/196030

**Authors:** Benjamin Rich Zendel, Greg West, Sylvie Belleville, Isabelle Peretz

**Author notes:** Corresponding author: Benjamin Zendel, Faculty of Medicine, Memorial University of Newfoundland, 230 Elizabeth Ave, St. John’s, NL, A1B 3X9, Canada.

## Abstract

It is well known that hearing abilities decline with age, and one of the most commonly reported hearing difficulties reported in older adults is a reduced ability to understand speech in noisy environments. Older musicians have an enhanced ability to understand speech in noise, and this has been associated with enhanced brain responses related to both speech processing and the deployment of attention, however the causal impact of music lessons in older adults is poorly understood. A sample of older adults was randomly assigned to learn to play piano (Mus), to learn to play a visuo-spatially demanding video-game (Vid), or to serve as a no-contact control (Nocon).After 6 months, the Mus group improved their ability to understand a word presented in loud background noise. This improvement was related to an earlier N100, enhanced P250 (P2/P3) and a reduced N600 (N400). These findings support the idea that music lessons provide a causal benefit to hearing abilities, and that this benefit is due to both enhanced encoding of speech stimuli, and enhanced deployment of attentional mechanisms towards the speech stimuli. Importantly, these findings suggest that music training could be used as a foundation to develop auditory rehabilitation programs for older adults.

## Introduction

Difficulties with hearing are one of the most commonly reported health issues in older adults, with more than 50% of people over age 60 reporting difficulties with hearing (Mathers, Smith, & Concha, 2001). Age-related decline in auditory perception can vary substantially between individuals and often includes difficulties understanding speech in adverse listening situations, such as when there is significant background noise (Pichora-Fuller, Schneider, & Daneman, 1995; Robert Frisina & Frisina, 1997; Schneider, Pichora-Fuller, & Daneman, 2010). These age-related changes in auditory perception are thought to reflect bilateral sensori-neural hearing loss due to physical changes in the inner ear (Gates & Mills, 2005; Stenklev & Laukli, 2004) as well as changes in the central auditory system (Alain, Snyder, & Dyson, 2006; Schneider et al., 2010). Difficulties with hearing have been associated with social isolation, and cognitive decline (Lin et al., 2013; Mick, Kawachi, & Lin, 2014). Given the prevalence, and negative outcomes of age-related decline in hearing abilities, finding ways to prevent, mitigate or delay these changes is of utmost importance, and evidence suggests that music training may be useful for preserving or enhancing auditory abilities in older adults. Here we report the results of a randomized control trial where one group of older participants received music training.

It is well known that musicians have enhanced auditory processing abilities (Kraus & Chandrasekaran, 2010), and these benefits are paralleled by an enhanced ability to understand speech in noisy environments (Parbery-Clark, Skoe, Lam, & Kraus, 2009; Zendel, Tremblay, Belleville, & Peretz, 2015). These cross-sectional studies implied that musical training causes neuroplasticity along the auditory pathway from the brainstem to structures throughout the cortex, and these brain changes support enhanced auditory abilities. Longitudinal research in younger adults confirmed that auditory benefits observed in musicians are due to neuroplasticity. Multiple studies with random assignment and control groups have demonstrated that after music training, participants have enhanced auditory abilities that are usually related to an enhanced neurophysiological measurement (Fujioka, Ross, Kakigi, Pantev, & Trainor, 2006; Kraus & White-Schwoch, 2015; Lappe, Herholz, Trainor, & Pantev, 2008; Tierney, Krizman, & Kraus, 2015). Emerging evidence suggests that enhanced auditory abilities persist into old age, with older musicians being able to understand speech in noisy environments better than older non-musicians (Parbery-Clark, Strait, Anderson, Hittner, & Kraus, 2011; Zendel & Alain, 2012). Moreover, longitudinal research using a non-music-based auditory training intervention, suggests that speech-in-noise perception can be improved in older adults (Anderson, White-Schwoch, Parbery-Clark, & Kraus, 2013). It remains unknown if music training in older adults can improve the ability to understand speech-in-noise.

Understanding speech in noise is a hierarchical process that occurs in multiple subcortical and cortical structures, and evidence suggests that musicianship and musical training can alter neural functions in multiple brain regions (Coffey, Mogilever, & Zatorre, 2017). In older musicians, there is evidence that enhanced endogenous, or attention-dependant processing contributes to their auditory benefit (Zendel & Alain, 2013, 2014). Benefits to subcortical processing of speech-in-noise in older adults have also been observed (Parbery-Clark, Anderson, Hittner, & Kraus, 2012); however these benefits seem to be reduced compared to the subcortical enhancements to speech-in-noise processing observed in younger musicians compared to non-musicians (Parbery-clark, Skoe, & Kraus, 2009). This pattern of results suggests that the musician benefit shifts from an exogenous processing benefit to an endogenous processing benefit as musicians age (Alain, Zendel, Hutka, & Bidelman, 2014).

Overall, these findings indicate that music training could be used in older adults as an engaging form of auditory training that could improve the ability to understand speech-in-noise. To determine if this is a possibility, we conducted a three-arm, single blind, randomized control study, where one group received music training. The ability to understand speech-in-noise and associated event-related brain responses were assessed at three time-points: before training, at the mid-point of training and after training. There are a number of ways to assess the ability to understand speech-in-noise, by varying both the target and the background noise. Different studies have used target stimuli that range from tones to speech phonemes to full sentences as a target, and, white noise, filtered white noise, and various forms of single-or multi-talker babble noise as the background noise (e.g. Billings, Tremblay, Stecker, & Tolin, 2009; Kaplan-Neeman, Kishon-Rabin, Henkin, & Muchnik, 2006; Martin, Sigal, Kurtzberg, & Stapells, 1997; Parbery-Clark et al., 2012; Pichora-Fuller, Schneider, & Daneman, 1995) To best understand if music training can improve the ability to understand speech in noise in the real world, we chose a paradigm that was ecological. This paradigm included real words as a target and multi-talker babble as the background noise. This paradigm was similar to one used previously, where younger musicians exhibited enhanced ability to understand words in noise compared to younger non-musicians (Zendel et al., 2015). Early event-related brain responses related to stimulus encoding were enhanced while later responses related to semantic processing were reduced in younger musicians compared to non-musicians (Zendel et al.2015). This suggests that musical training enhances the representation of an incoming speech stimulus, which facilitates later semantic access. This paradigm is therefore well suited to examine the impact of music lessons as it is sensitive to musicianship for both behavioural and neurophysiological measures, and it is based on natural speech sounds.

## Methods

### Design

The study was designed as a three-arm single blind randomized control trial. Participants were randomized into three groups using a stratified covariate-adaptive procedure (see below). Participants took part in three testing sessions. The Pre-training session (Pre) took place before the intervention, the Mid-training session (Mid) took place 3 months after the start of the intervention, and the Post-training session (Post) took place 6 months after the onset of the intervention. During the Pre, Mid and Post sessions, participants completed a series of auditory and cognitive assessments. Here we report the results of a speech-in-noise task that was done while monitoring electrical brain activity (EEG). The results of other assessments will be presented elsewhere.

### Participants & Randomization Procedure

Participants were recruited into the study from the Centre de Recherche, Institut Universitaire de Gériatrie de Montréal participant pool. The study received ethical approval from the Comite conjoint d’evaluation scientifique – Regroupment Neuroimagerie/Quebec (CES-RNQ).

Participants were pre-screened to ensure that they did not have any present or past major illness, did not meet criteria for Mild Cognitive Impairment (MCI) using the Montreal Cognitive Assessment (Nasreddine et al., 2005), were not taking any psychiatric medications or medication known to have an impact on cognition, were MRI compatible, were a non-video game player and a non-musician. To be considered a non-musician, participants had to not currently play a musical instrument, and had no more than 3 years of formal music training in their life. Music lessons that were part of the normal education curriculum were not included. To be considered a non-video game player, participants had little to no experience with commercial video games (e.g., games played on a computer or game console) during their lifetime. Casual games such as computerized card or puzzle games were not considered to be video games.

All participants were randomized into one of three groups. Randomization was done by an independent research assistant, using a predefined randomization table prior to contacting participants to ensure that participants were blind to the existence of the other two groups. Randomization was stratified using a covariate-adaptive randomization procedure. Each factor was stratified into two categories. For the factor of age there was “younger” (55-64 yrs) and “older” (65-75 yrs); for the factor of education there was low (< 16 yrs) and high (> 16 yrs); and for the factor of gender there was female and male. Because participants were recruited from a database, age, education level, and gender of each participant were known before they were contacted and it was thus possible to stratify randomization on the basis of these three factors. This stratification led to eight possible stratification groups. Lists of participants were provided from the participant database to the research assistant. Based on the stratification variables, the participant was assigned to one of the eight stratification groups based on the demographics available from the participant database (e.g. female, younger, high education; male, older, low education; etc…). Each of these eight groups was assigned a random but balanced order to determine experimental group assignment. That is the first person contacted who was in the “female, younger, higher education” stratification group was invited to participate in the experimental Music Training group (Mus; see below for details). If this participant accepted she became participant 1 in the Mus group. If she refused, the next person contacted in the “female, younger, higher education” stratification group was invited to participate in the Mus group. This repeated until a person in this stratification group volunteered to participate in the Mus group. Next, people in the same stratification group were invited to participate in the No Contact Control group (Nocon; see below for details) until one person volunteered to participate. Finally, a person in this stratification group was invited to participate in the Video Game training group (Vid; see below for details) until one person volunteered to participate. This procedure was repeated, except the order of recruitment in experimental group was randomized for each cycle of three assessments. That is each three participants were recruited into one of the three groups (Music, Video, Control), but the order in which they were recruited was random. The orders were also randomized across all the stratification groups. Accordingly, participants who choose not to participate were not included in the randomization matrix.

Forty-five participants in total were recruited into the study. Using the stratified randomization procedure, 15 participants were assigned to the Vid group, 15 participants were assigned to the Mus group and 15 participants were assigned to the Nocon group. During the study, 2 participants withdrew from the Mus group, 2 withdrew from the control group, while 11 withdrew from the Vid group. To compensate for the higher attrition rate within the Vid group, an additional four participants were assigned who were matched for the age, gender and education of the other two groups, however, the stratified randomization procedure was not used. This resulted in a total of 8 participants completing the training within the Vid group. The demographics of the participants within each group are presented in Table 1. Due to a participants’ hair style, EEG data was unusable from one participant in the Mus group, however this participant still completed the behavioural task. For more details about the withdrawal rate from the Vid group see West, Zendel, Konishi, Benady-Chorney, Bohbot, Peretz & Belleville (submitted).

**Table 1:** Participant demographics

| Group | Age (SD) | Gender | Yrs. Education (SD) |
| --- | --- | --- | --- |
| **Music**<br>**(n = 13)** | 67.5 (4.2) | 10 female;<br>3 male | 14.5 (2.2) |
| **Video**<br>**(n = 8)** | 66.9 (3.9) | 4 female;<br>4 male | 17.5 (2.3) |
| **No**<br>**Contact**<br>**(n = 13)** | 69.3 (5.7) | 10 female;<br>3 male | 15.2 (3.1) |

### Training Procedure

#### *Piano training* group (Mus)

Piano training was done at home using Synthesia software, and an 88-key M-Audio MIDI piano. Synthesia is a piece of software that uses a non-standard form of musical notation that can be understood within a few minutes. This was critical as learning to read traditional music notation can take a long time. Notes in Synthesia are presented as coloured bars that fall from the top of the computer screen. At the bottom of the screen is an image of a piano keyboard, and when a coloured bar hits a certain note, the participant plays that note. The length of the bar indicates how long to hold the note for. First, a research assistant installed and calibrated the piano to work on the participant’s home computer. Next, the participant completed an introductory lesson that included introductory information about music, detailed instructions on how to use Synthesia, and directions on how to record their progress. Introductory music information included lessons about note names, how to place hands on the piano, and how to synchronize performance with the information on the screen and the metronome. A set of introductory lessons, and beginner piano music was installed on the computer. Participants were told to start with the lessons, and once they were comfortable with the lessons, to try out some of the introductory songs. Participants were encouraged to move at their own pace, but to try to master a given lesson or song before moving on. Sometimes participants would work on a lesson and song simultaneously. The goal was to keep participants as engaged as possible in the piano lessons.

#### *Video game training group* (Vid)

Video game training was done at home using the Nintendo Wii console system equipped with a Wii Classic Controller. All participants in this group trained on Super Mario 64. Two participants completed all task within Super Mario 64 before the completion of the 6-month training period. In these cases, they continued to train on a very similar game, Super Mario Galaxy, until the end of the training period. Super Mario 64 and Super Mario Galaxy are three-dimensional platform games where the player is tasked with exploring a virtual environment to search for stars (tokens). When enough stars are collected through completing in-game goals, the player can then progress further into the game and will encounter new environments to explore.

After the participant completed the pre-tests, a research assistant installed the Nintendo Wii to the participant’s home television. The research assistant then gave an initial orientation to the participant to teach them how to turn on the Nintendo Wii and access the Super Mario 64 game. This was followed by a custom in-game orientation which taught the participant to move the character around the virtual environment. At this point, some participants encountered certain challenges associated with maneuvering the character. Some had issues with understanding the game’s mechanics. Further, Super Mario 64 has a very steep learning curve that was not originally designed to be played by someone with little to no video game or computer experience. For this reason, the research assistant returned to the participant’s home for up to three additional supervised 2-hour training sessions to teach the participant how to properly maneuver the character and progress through the game. After this, participants were given a custom made instruction booklet which outlined how and where to collect all the stars for the first four levels. This allowed participants to learn the game’s mechanics in further detail and practice the basic motor coordination that was required. After this point, participants had to search for and obtain the stars within each remaining level without any assistance from the research team.

#### No-contact Control Group (Nocon)

The no contanct control group had no contact with the research team during the six-month period other than to complete the pre-training, mid-term and post-testing sessions.

Music and Video game training lasted six months. In all cases, participants kept a record of their daily training progress and were asked to complete a minimum of 30 minutes of training at least five days a week, although some completed more than this amount. All participants were told that they were expected to improve in performance. Participants in the Vid group were told that there was evidence that video game training enhances cognitive abilities, and that video game training in older adults was expected to improve those abilities. Participants in the Mus group were told that there was evidence that musicians have enhanced cognitive abilities, and that we expected musical training to improve those abilities. Finally, the Nocon group was told that we were investigating test-retest effects, and that they were expected to improve on the experimental tasks. All participants were debriefed about the other groups at the end of the final testing session.

### Stimuli

Stimuli were 150 French words spoken by a male, from a list of phonetically balanced, equally understandable monosyllabic words (see Picard 1984). Words were presented at ∼75 decibels sound pressure level (dB SPL), through insert-earphones (Etymotic ER-2), as measured by a sound level meter (Quest Technologies) that measured the amplitude of the stimuli presented from the one insert earphone. Words were presented in three conditions. In one condition the words were presented in isolation (None). In the other two of the conditions, multi-talker babble noise was presented with the words at ∼60 dB SPL (Quiet noise; 15 dB signal-to-noise ratio [SNR]) and ∼75 dB SPL (Loud noise; 0 dB SNR). The multi-talker babble was created by individually recording four native speakers of French (two female, two male), each reading a rehearsed monologue in a sound-attenuated room for 10 minutes. The recordings were made at a sampling rate of 44.1 KHz at 16 bits, using an Audio-Technica 4040 condenser microphone. The individual recordings of each monologue were normalized, and combined into a single monaural sound file using Adobe Audition (Version 10). The 10-minute multi-talker babble noise was looped repeatedly during listening conditions where the multi-talker babble was present.

### Procedure

The words were presented in a random order, in three levels of multi-talker babble noise. In the ‘None’ condition, words were presented without multi-talker babble noise. In the Quiet noise condition words were presented with multi-talker babble noise that was 15 dB below the level of word (i.e., 15 dB signal-to-noise ratio [SNR]), while in the Loud-noise condition words were presented with multi-talker babble noise that was at the same level as the word (i.e., 0 dB SNR). In addition, all three noise-levels were presented in two listening conditions, Active and Passive. In the passive condition, participants were told to ignore the words and watched a self-selected silent subtitled movie. Words were presented with a stimulus onset asynchrony that was randomized between 2500-3500 ms. The use of muted subtitled movies has been shown to effectively capture attention without interfering with auditory processing (Pettigrew at al., 2004). In the active condition, participants were told to repeat the word aloud. To avoid muscle artifacts in the ERPs, participants were told to delay their response until they saw a small LED light flash 2000 ms after the presentation of the word. Word correctness was judged online by a native French speaker. The research assistant performing the judgment had the text of word presented in front of them on a screen, and did not hear background noise during the word repetition. The research assistant was told to score the word as correct if it was understandable as the word written on the screen. If there was confusion, the research assistant asked the participant to repeat the word until the research assistant was sure the repetition was a match or not. We chose to use word repetitions because it required an accurate lexical match of the incoming word in order to correctly repeat it back, and pilot testing confirmed that the delayed oral response did not contaminate the ERPs with muscle artifacts. An alternative, would have been to use a forced choice procedure, however this would likely create a biased estimate of word understanding because the presentation of choices limits what a participant can report, and may bias their performance if they were able to hear part of the word.

### Recording and averaging of electrical brain activity

Neuroelectric brain activity was digitized continuously from 70 active electrodes at a sampling rate of 1024 Hz, using a Biosemi ActiveTwo system (Biosemi, Inc., Netherlands). Six electrodes were placed bilaterally at mastoid, inferior ocular, and lateral ocular sites (M1, M2, IO1, IO2, LO1, LO2). All averages were computed using Brain Electrical Source Analysis (BESA) software, version 6.1. The analysis epoch included 200 milliseconds of pre-stimulus activity and 1500 milliseconds of post-stimulus activity. Trials containing excessive noise (> 120 μV) at electrodes not adjacent to the eyes (i.e., IO1, IO2, LO1, LO2, FP1, FP2, FPz, FP9, and FP10) were rejected before averaging. Continuous EEG was then averaged separately for each condition, into six ERPs; Active: None, Quiet & Loud, and Passive: None, Quiet, and Loud, at each electrode site. Prototypical eye blinks and eye movements were recorded before the start of the study. A principal component analysis of these averaged recordings provided a set of components that best explained the eye movements. These components were then decomposed into a linear combination along with topographical components that reflect brain activity. This linear combination allowed the scalp projections of the artifact components to be subtracted from the experimental ERPs to minimize ocular contamination such as blinks, vertical, and lateral eye movements for each individual average with minimal effects on brain activity (Berg & Scherg, 1994). After this correction, trials with greater than 120 μV of activity were considered artifacts, and excluded from further analysis. In addition, during active listening, trials where the participant did not correctly repeat the word were excluded from the analysis. To determine if the experimental manipulation had an impact on the number of trials accepted, a mixed design ANOVA was calculated that included Session, Noise Level, and Listening Condition as within subject factors, and Group as a between subject factor. As expected, there were significant main effects of both Noise Level, F (2, 60) = 55.79, p <.001 & Listening Condition, F (1, 30) = 85.0, p <.001. Additionally, there was a significant interaction between Noise Level and Listening Condition F (2, 60) = 36.2, p <.001. No other effects or interactions were significant. The ERP analysis included an average of 121, 118, and 90 trials during the None, Quiet and Loud Noise Level, respectively, during Active listening, and 135, 134, and 131 trials during the None, Quiet and Loud Noise Level during Passive listening. This means that there were fewer observations during the Active Listening condition, and fewer still during the Loud Noise condition during Active listening. The advantage to this approach was that the brain activity during Active trials only included successful understanding of the speech. This permits us to connect brain activity to understanding. The disadvantage of this approach is that the individual averaged ERPs from the Loud Active trials are likely more variable. By using a pre-post design, and comparing differences in the same condition, the effect of this variability is minimized. Finally, ERPs were band-pass filtered to attenuate frequencies below 0.1 Hz, and above 15 Hz, and referenced to the linked mastoid.

## Results

Given that this study was designed as an RCT, the critical effect in all the data analysis will be based on Group by Session interactions, followed-up by a significant effect in the Mus Group, and non-significant effects in the other groups. In order to fully explore the data, other effects are reported as well.

### Behavioural data

Data was analyzed using a mixed design ANOVA that included Session (Pre, Mid, Post) and Noise level (None, Quiet, Loud) as within subject factors and Group (Mus, Vid, Nocon) as a between subject factor. There was no behavioural data collected during passive trials, so Listening Condition was not included as a factor. Accuracy was impacted by Noise Level, F (2, 62) = 263.63, p <.001, η^2^ =.9, with accuracy being higher in the None condition compared to Quiet noise (p <.001), and Quiet noise being higher than Loud (p <.001; see Figure 1). The three-way Noise level by Session by Group was nearly significant, F (8, 124) = 1.91, p =.064, η^2^ =.11. Follow-up simple two-way ANOVAs revealed a significant Noise level by Session interaction for the Mus group, F (4, 48) = 3.25, p =.019, η^2^ =.21, but not for the Vid or Nocon groups (p =.90 &.16, respectively) demonstrating that impact of Session was only significant for the Mus group. Further follow-up tests in the Mus group revealed that accuracy improved in the Loud condition, F (2, 24) = 3.99, p =.032, η^2^ =.25, but not in the Quiet or None conditions (p =.45 &.88, respectively). To ensure groups were balanced at baseline, a series of one-way between subject ANOVAs were done for each Noise level. None of these effects were significant (p =.45,.84 &.56 for Loud, Quiet and None, respectively).

**Figure 1:**
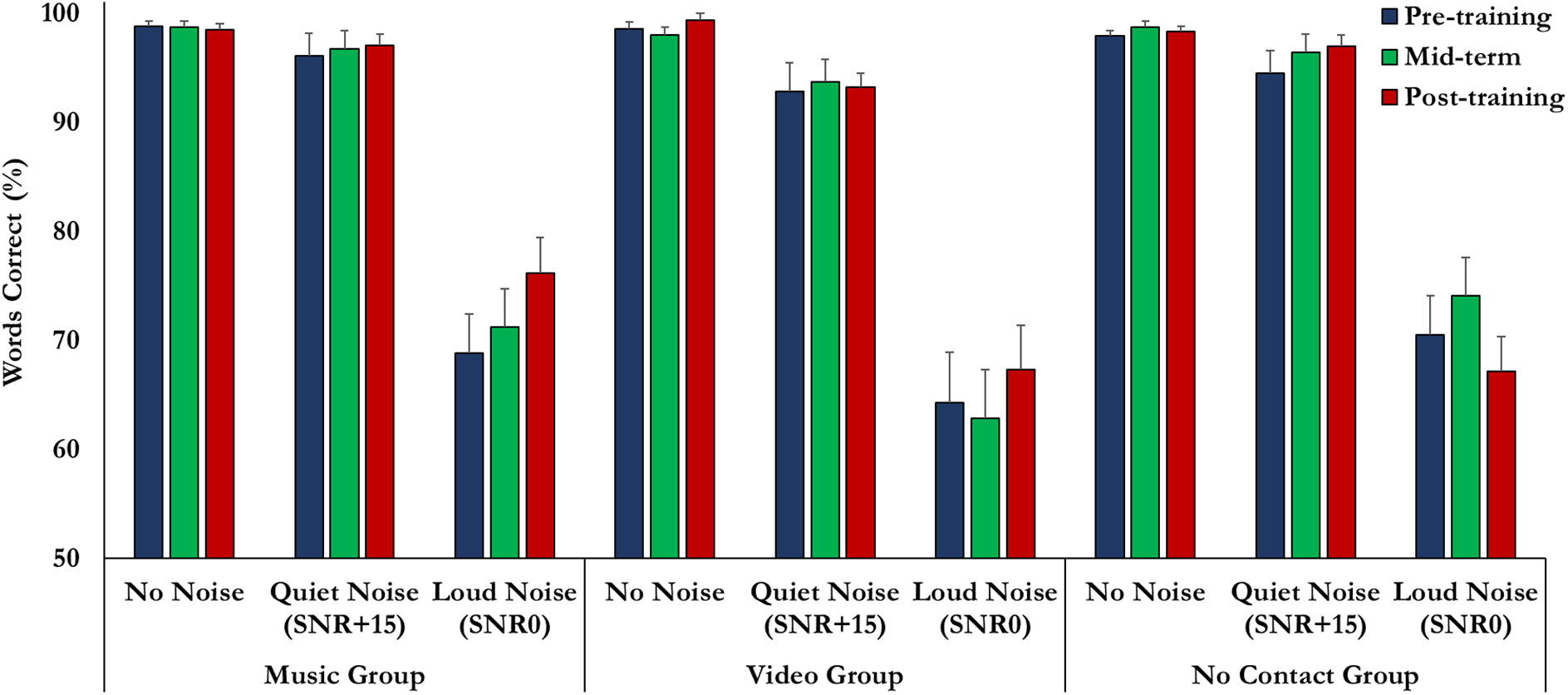
Accuracy during the active listening task. Participants in all three groups were able to repeat nearly all the words accurately during the None and Quiet Noise conditions. Accuracy improved during the Loud Noise condition in the Mus group from pre-training to post-training. No improvement was observed in either of the other groups.

### ERP Results

The ERP analysis focused on the first three deflections that appeared to be present in all three noise levels during active listening. The components are named based on their latency during the None Noise level condition during Active Listening. Accordingly, we focused on an N100, a P250 and an N600.

As a first step, we wanted to determine if there were any differences between the two control groups (Vid & Nocon) on the three ERP components. We found no differences between these groups, thus to improve statistical power, we include them as a single Control group (Con). All EEG data was analyzed using Session (Pre, Mid, Post), Listening Condition (Active, Passive), Noise Level (None, Quiet, Loud) and Electrode (different montages were used for different components) as within-subject factors and Group (Mus, Con) as between subject factors. Interactions and main effects of electrode are not reported as multiple electrodes were used to gain a stable and reliable estimate of the relevant component.

### ERP: N100

Peak N100 amplitude and latency were extracted from a montage of fronto-central electrodes (F1, Fz, F2, FC1, FCz, FC2, C1, Cz, C2). A long extraction window was used because of the impact of Noise level on ERP latencies (90-390 ms). The N100 can be seen in Figure 2A.

**Figure 2:**
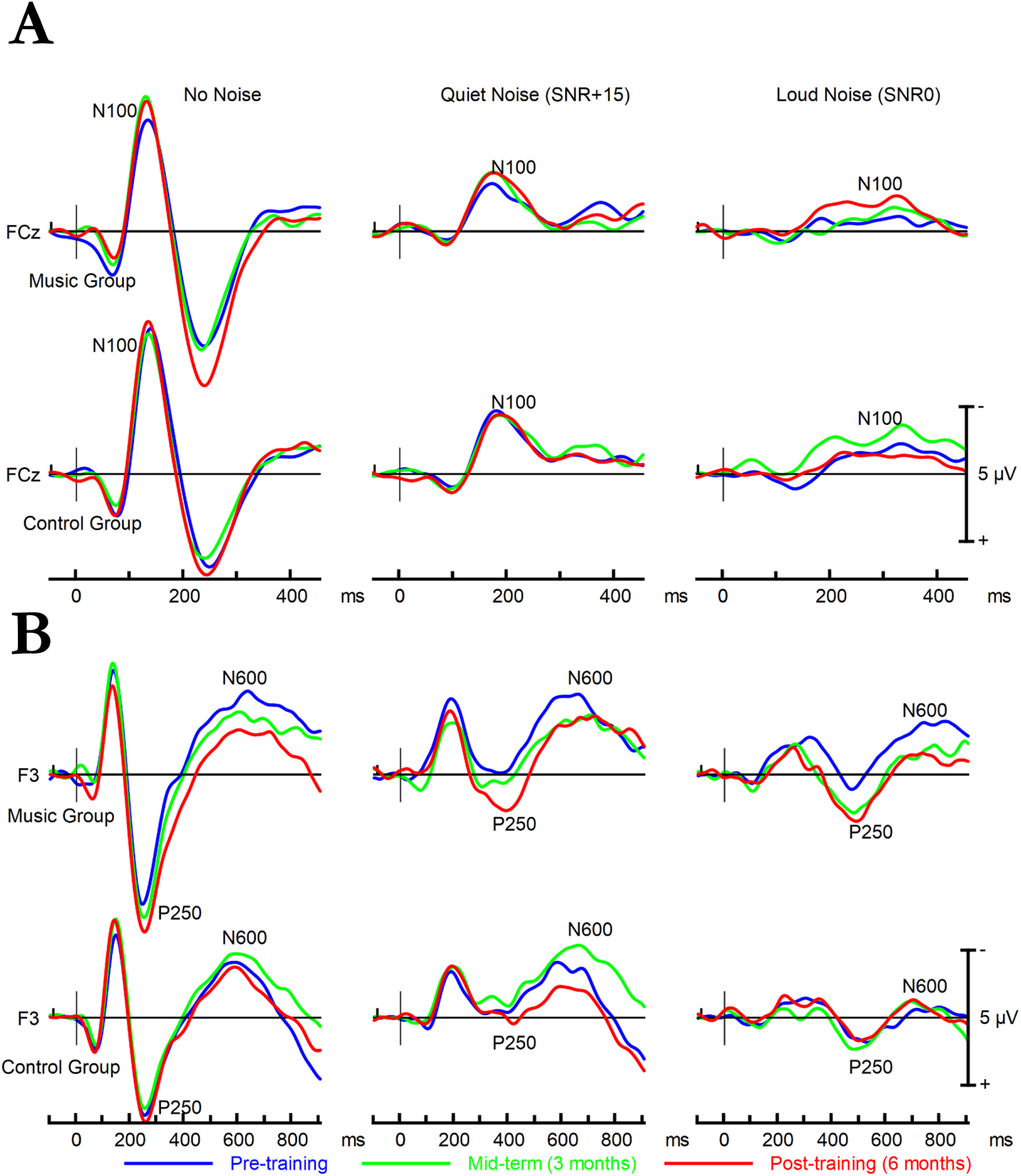
(A) ERP plots from electrode FCz recorded during passive listening. The N100 increased in amplitude in the Mus group across all noise levels from pre-training to post-training. No change in N100 amplitude was observed in the other groups. N100 is identified on all plots. As noise level increased the N100 was delayed and reduced in amplitude. (B) ERP plots from electrode F3 recorded during active listening. The P250 increased in amplitude in the Music group across all noise levels from pre-training to post-training. No change in P250 was observed in the Combined Control group. The P250 is identified on all plots. In the None condition the response appears like a P2, however, during the Quiet and Loud Noise conditions it is delayed and appears more like a P3-type response. The N600 decreased in amplitude in the Mus group across all noise levels from pre-training to post-training. No change in N600 was observed in the other groups. The N600 is identified on all plots.

### N100 Amplitude

Overall N100 amplitude was impacted by Noise Level during both Active and Passive listening, F (2, 62) = 53.6, p <.001, η^2^ =.63; polynomial decompositions revealed that N100 amplitude decreased as noise level increased, F (1, 31) = 60.14, p <.001, η^2^ =.66. The impact of Noise Level on N100 amplitude was qualified by Session as the Noise Level by Session interaction was significant, F (4, 124) = 2.59, p =.04, η^2^ =.08, and this effect was similar across all groups as the Noise Level by Session by Group interaction was not significant (p =.43). Follow-up tests revealed a significant effect of Noise Level during all three Sessions F (2, 62) = 24.95, p <.001, η^2^ =.44; F (2, 62) = 52.70, p <.001, η^2^ =.62; F (2, 62) = 41.59, p <.001, η^2^ =.57; (Pre, Mid, Post, respectively). The difference between each noise level during each session was significant (p <.001), except for the difference between Quiet and Loud Noise during the Mid session (p =.047), and the difference between Quiet and Loud Noise during the Post Session (p =.01). These small, but statistically significant differences are the likely source of the Noise Level by Session interaction across groups and listening conditions.

### N100 Amplitude: effect of training

Most importantly, the Group by Session by Listening Condition interaction was significant, F (2, 62)= 3.28, p =.044, η^2^ =.10. Follow-up test revealed that the Session by Listening condition interaction was significant in the Mus group, F (2,22) = 4.75, p =.019, η^2^ =.30. The Session by Listening Condition interaction and main effect of Session were not significant in the Con group (p=.50,.37, respectively). In the Mus group polynomial decompositions were calculated to model the linear trend of the N100 amplitude across testing sessions. These revealed that N100 amplitude increased linearly from the Pre to Mid to Post Session during Passive listening, F (1, 11) = 5.43, p =.04, η^2^ =.33, but not Active listening (p =.70).

### N100 Latency

Overall N100 latency was impacted by Noise Level, F (2, 62) = 178.71, p <.001, η^2^ =.85; polynomial decompositions revealed that N100 latency increased linearly as Noise Level increased, F (1, 31) = 260.73, p <.001, η^2^ =.89. The effect of Noise Level was impacted by Listening Condition, F (2, 62) = 3.52, p =.036, η^2^ =.10. Follow-up tests revealed that N100 Latency was similar during Active and Passive listening when Noise Level was None or Quiet (p =.18 &.29); however, N100 Latency was longer during Passive Listening when Noise Level was Loud F (1, 32) = 3.8, p =.06, η^2^ =.11.

### N100 Latency: effect of training

Most importantly, the Session by Group interaction was significant, F (2, 62) = 5.40, p =.007, η^2^ =.15. Follow-up pairwise comparisons revealed that N100 latency in the Mus group decreased from Pre to Mid (p =.016) and was stable between the Mid and Post Sessions (p =.16). In the Con group, N100 Latency was similar in the Pre and Mid Session (p =.18), and the Pre and Post session (p =.30).

### ERP: P250

Peak P250 amplitude and latency were extracted from a montage of fronto-left electrodes (F1, F3, F5, F7, FC1, FC3, FC5, FC7) during the 200-600 ms epoch. The P250 can be seen in Figure 2B. A detailed discussion about whether this component is a P2 or P3 is included in the discussion.

### P250 Amplitude

Overall, P250 was larger during active listening compared to passive listening F (1, 31) = 7.44, p =.01, η^2^ =.19. The P250 was impacted by noise level F (2, 62) = 90.49, p <.001, η^2^ =.75, and pairwise comparisons revealed that P3 was smaller in the Quiet condition compared to both the None and Loud conditions (p <.001 for both). The Listening Condition by Noise Level interaction was significant F (2, 62) = 3.94, p =.025 η^2^ =.11. Follow-up tests revealed that P250 amplitude was larger during Active listening when there was no background noise, t (32) = 2.96, p =.006, η^2^ =.22, and when there was loud background noise t (32) = 2.68, p =.012, η^2^ =.18. P3 amplitude was not different between active and passive listening when there was Quiet background noise (p =.6)

### P250 Amplitude: effect of training

Most importantly, the Group by Session by Listening Condition interaction was significant, F (2, 62) = 3.79, p =.028, η^2^ =.11. Follow-up tests revealed a significant interaction between Listening Condition and Session in the Mus group, F (2, 22) = 3.95, p =.034, η^2^ =.26. This interaction and the main effect of session were not significant in the Con group (p =.31 & 30, respectively).Follow-up tests in the Mus group revealed a significant effect of Session during Active Listening F (2, 22) = 3.54, p =.047, η^2^ =.24. Polynomial decompositions further revealed that P250 amplitude increased during active listening in a linear manner from Pre to Mid to Post, F (1, 11) = 4.72, p =.053, η^2^ =.30. The effect of Session on P250 amplitude was not significant in the Mus group during Passive listening, F (2, 22) = 0.35, p =.71, η^2^ =.03.

### P250 Latency

P250 Latency was not impacted by training, as the main effect of Session (p =.46) and all interactions with Session were not significant (all p >.19). P250 latency increased as the background noise level increased, F (2, 62) = 119.06, p <.001, η^2^ =.79, however this effect was impacted by Listening condition as the Listening Condition by Noise level Interaction was significant F (2, 62) = 3.98, p =.024, η^2^ =.11. There was no difference between active and passive listening for P250 latency during the None condition (p =.26) nor when background noise was loud (p =.13). The P250 was earlier during active listening compared to passive listening when there was Quiet Noise, t (32) = 2.14, p =.04, η^2^ =.13.

### ERP: N600

Peak N600 amplitude and latency were extracted from a montage of fronto-left electrodes (F1, F3, F5, F7, FC1, FC3, FC5, FC7) during the 400-900 ms epoch. The N600 can be seen in Figure 2B.

### N600 Amplitude

Overall N600 amplitude was larger during active listening compared to passive listening, F (1, 31) = 32.1, p <.001, η^2^ =.51. N600 amplitude was also impacted by Noise Level F (2, 62) = 18.2, p <.001, η^2^ =.37, and was largest during the Quiet condition compared to both None and Loud (p <.01 for both). These two effects were qualified by a significant interaction between Listening Condition and Noise Level F (2, 62) = 5.03, p =.009, η^2^ =.14. Follow-up tests revealed a significant effect of Noise Level during Active Listening, F (2, 62) = 14.95, p <.001, η^2^ =.32, but not during Passive Listening (p =.84). During Active Listening N600 amplitude was largest when Noise Level was Quiet compared to both None and Loud (p <.001 for both).

### N600 Amplitude: effect of training

More importantly, the Session by Listening Condition by Group interaction was significant F (2, 62) = 4.37, p =.017, η^2^ =.14. Follow-up tests revealed that the Session by Listening Condition interaction was significant in the Mus group F (2, 22) = 5.44, p =.012, η^2^ =.33. This interaction and the main effect of Session were not significant in the Con group (p =.38 &.45, respectively).Further follow-up tests in the Mus group revealed that the effect of Session was significant during Active Listening, F (2, 22) = 3.57, p =.045, η^2^ =.25, and significant at a trend level during Passive Listening (p =.07). In the Music group during active listening, N600 amplitude decreased from Pre to Mid (p =.046), but was stable between Mid and Post (p =.76). Interestingly, the amplitude of the N600 increased from Pre to Post in the Music group during Passive listening (p =.038).

### N600 Latency

Overall N600 Latency was earlier during passive listening, F (1, 31) = 24.76, p <.001, η^2^ =.44. The interaction between Listening Condition and Noise Level was significant, F (2, 62) = 8.61, p =.001, η^2^ =.22. Follow-up tests revealed that N600 Latency was impacted by Noise Level during Active Listening, F (2, 62) = 7.06, p =.002, η^2^ =.18, but not Passive Listening (P =.12). During Active Listening, N600 Latency increased from None to Quiet (p =.004) and from None to Loud (p =.002).

### N600 Amplitude: effect of training

More importantly, the Session by Listening Condition by Group was significant, F (2, 62) = 4.90, p =.011, η^2^ =.14. Follow-up tests revealed that the Session by Listening condition was significant in the Mus group, F (2, 22) = 5.96 p =.009, η^2^ =.35, but not the Con group (p =.58). Follow-up tests in the Mus group revealed a significant effect of Session during Passive Listening F (2, 22) = 8.86, p =.002, η^2^ =.45, but not Active Listening (p =.63), with N600 latency increasing from Pre to Post (p =.002)

The Noise Level by Group interaction was also significant, F (2, 62) = 3.39, p =.04, η^2^ =.10. Follow-up tests revealed that the effect of Noise Level was significant in the Mus group F (2, 22) = 3.53, p =.047, η^2^ =.24, as N600 latency was longer when Noise Level was Loud compared to None (p =.036). There was no effect of Noise level in the Con group (p =.65). This effect was consistent across Sessions, as the Noise Level by Session by Group interaction was not significant (p =.23).

### Brain-Behaviour Pre-Post Difference Correlations

The next step in the analysis was to determine if the training-related change in neurophysiological response to a word in noise predicted the behavioural change. Significant Group by Session effects were observed for N100 latency, P250 amplitude and the N600 amplitude during Active listening. Accordingly, we focused on N100 latency, P250 amplitude and N600 amplitude. For all three electrophysiological measurements, data was averaged across the analysis montage reported above for the pre-and post-training conditions (i.e., N100: nine fronto-cental electrodes, P250 & N600: eight fronto-left electrodes). The difference was calculated between the pre-and post-training conditions for the three electrophysiological measurements and the for accuracy. Given that accuracy was near ceiling in both the No-noise and Quiet-noise conditions, the analysis focused on data from the Loud-noise listening condition only. There was a trend towards the decrease in N100 latency being associated with improved performance, r (33) = -.32, p =.074. Interestingly, the strength of this correlation may have been larger in the Mus Group r (12) = -.38, p =.23, compared to the Con Group, r (21) = -.28, p =.22. Participants who exhibited the most improvement in accuracy also had the largest gain in P250 amplitude, r (33) =.44, p =.01. Interestingly, this correlation was similar in both the Mus Group r (12) =.53, p =.08, and in the Con Group, r (21) =.60, p =.004. Participants who exhibited the most improvement in accuracy also had the largest decrease in N600 amplitude, r (33) =.54, p =.001. As before, this correlation was similar in both the Mus Group r (12) =.54, p =.07, and in the Con Group, r (21) =.59, p =.005. Note that a larger N600 amplitude is represented by a smaller number (i.e., more negative), thus a positive correlation indicates that when accuracy increased, N600 decreased.

Given that the correlations for the pre-post difference in N100 latency, P250 and N600 with the pre-post difference in behaviour were similar during the Loud Active condition, we calculated a regression with the pre-post difference in behaviour performance entered as a dependant variable, and the pre-post difference in N100 latency, P250 and N600 entered as predictor variables. The overall regression model was significant, F (3, 29) = 4.19, p =.014, R^2^ =.30, while none of the individual factors independently predicted a significant portion of the variance in the pre-post difference in behaviour, N1lat: β = -.12, p =.48; P3: β =.07, p =.78; N600: β =.44, p =.09. At the same time, bivariate correlations between the pre-post difference in N100 latency and P250, N100 latency and N600, and P250 and N600 were all significant r(33) = -.33, -.39 &.77, p =.06,.01 & <.001, respectively. This pattern suggests multicolliniarity, therefore, a principal component analysis (PCA) was computed for the pre-post difference in N100 latency, P250 amplitude and N600 amplitude. The PCA revealed a single underlying component that accounted for 67.5% of the variance. This component predicted the pre-post change in accuracy during the Loud-noise condition, r (33) =.54, p =.001 (Figure 3)

**Figure 3:**
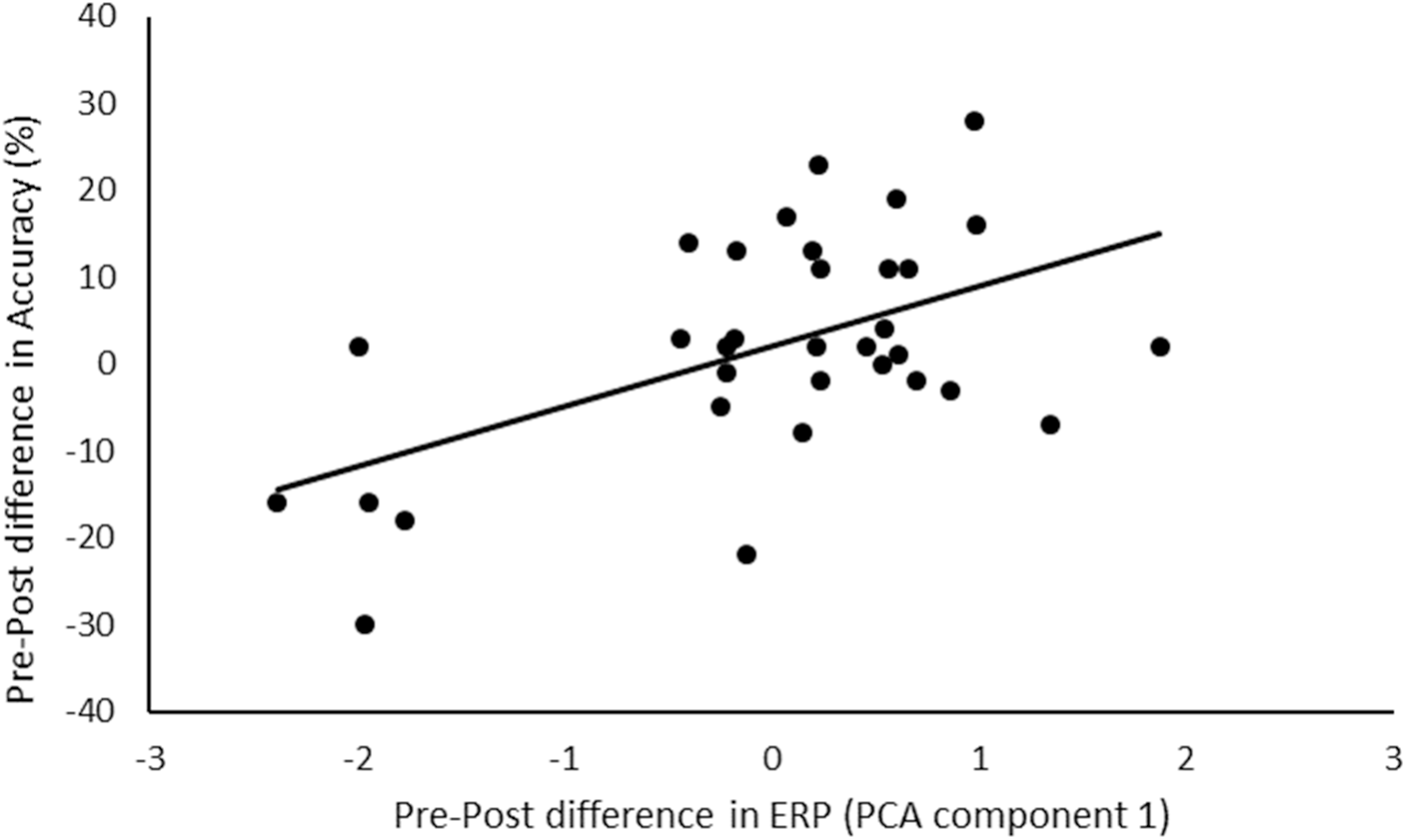
Post minus Pre difference in accuracy as a function of the first PCA component extracted from the post minus pre difference for N100, N250 and N600. This component accounts for 60.7% of the variance in the post minus pre ERP data.

## Discussion

Six months of self-directed music lessons improved the ability to understand speech in background noise. This was related to decreased N100 latency, increased P250 amplitude and decreased N600 amplitude. Training-related change to N100 latency, P250 and N600, evoked when background noise was the loudest predicted improvement in the ability to understand speech in background noise. There was also a post-training enhancement in the Mus group to the N100 response during passive listening. Overall, these results suggest that music training can be used to improve the ability to understand speech in noise in older adults by improving how older adults process speech and deploy their attentional resources to speech stimuli.

The most critical finding from this study is that music lessons improved the ability to understand speech in noise for older adults. This finding is critical for at least two reasons. The first is that it demonstrates that music training has a causal impact on hearing abilities in older adults. This provides support for previous work that demonstrated that older musicians have an advantage in understanding speech-in-noise (Zendel et al. 2012; Parbery-Clark et al. 2011). The second reason this finding is critical is that it highlights that the brain remains plastic in older adults and is susceptible to be modified based on experiences. In the current context this is critical, as hearing difficulties in older adults are nearly universal (Gates & Mills, 2005; Quaranta et al., 2015), and one of the most commonly reported hearing difficulty is understanding speech in noisy environments. Accordingly, the results of this study demonstrate that auditory rehabilitation is likely to be successful in older adults, and that using music-lessons as a model for the rehabilitation is a good starting point.

Another critical component of this study is that it identified the neural mechanisms associated with improved abilities to understand speech-in-noise. The earliest benefit of music training was observed on the auditory N100. Music training hastened the N100 across all listening conditions, and enhanced it during passive listening. The enhanced and earlier N100 suggests that music lessons can improve the encoding of incoming acoustic information. It is well established that the N100 is associated with the physical properties of the stimulus (Näätänen & Picton, 1987). Given that the noise was continuous during the presentation of stimuli, an earlier N100 suggests that after music training, the ascending auditory pathway processed a target speech input faster, regardless of noise level. Moreover, this speeded processing was related to improved ability to understand speech-in-noise. This provides support for the idea that musical training leads to more robust encoding of auditory features (Parbery-Clark et al., 2012). Interestingly, the enhancement to N100 was only observed during passive listening. The lack of enhancement during active listening may be due to the engagement of an attentional processes, that was associated with an increase in positive going electrical brain activity over frontal electrodes. Indeed, the positive going component following the N100 (See *P250: P2 or P3?* below) was enhanced after training during active listening. It is likely that the earlier N100 after training allows for attentional processes to engage earlier as the pre-post difference in N100 Latency was related to the enhancement of the P250. Overall, this suggests that music training improved the encoding of basic auditory features, reflected in an earlier and larger N100. This enhanced encoding would have implications for later stages of processing related to orienting attention and extracting meaning from the auditory stimulus.

## P250: P2 or P3?

Before the impact of music training on the attention dependant ERP components can be discussed, the impact of noise on the latency of the evoked responses should be addressed. One difficulty interpreting the current results is that the second positive peak could be considered a P2 or P3 depending on the noise level. When measuring cortical responses to stimuli in background the N1-P2 complex becomes reduced in amplitude and delayed as the noise level increased (Billings et al., 2009; Kaplan-Neeman et al., 2006; Martin et al., 1997; Zendel et al., 2015). Not surprisingly, we observed the same effect in the current study. One of the effects of music training was on the second positive component at fronto-left electrodes. This component was likely a P2 during the no-noise condition (peak ∼250 ms), but was delayed by the addition of noise such that during the loud-noise condition the peak was closer to 500 ms, and thus more like a P3-type response. The P2 response is generally associated with stimulus classification (Crowley & Colrain, 2004), and is enhanced in younger musicians, particularly as the stimulus increases in spectral complexity (Shahin, Bosnyak, Trainor, & Roberts, 2003; Shahin, Roberts, Pantev, Trainor, & Ross, 2005). Critically, the P2 is an obligatory response that will occur regardless of where the listeners attention is focused (Crowley & Colrain, 2004), while P3 responses are typically related to allocation of cognitive resources for conscious information processing (Dinteren, Arns, Jongsma, & Kessels, 2014; Kok, 2001).

Typically, P3 responses decline in older adults, reflecting an age-related decline in allocation of attention (Amenedo & Díaz, 1998; Anderer, Semlitsch, & Saletu, 1996; Dinteren et al., 2014). Moreover, an anterior shift is often observed in older adults when processing sensory information (Davis, Dennis, Daselaar, Fleck, & Cabeza, 2008; Haxby & Maisog, 1994). This frontal shift is likely a compensatory mechanism that offsets age-related decline in functional and structural properties of posterior brain regions. In the current context, the training-related enhanced P250 was observed over frontal-left regions, during active listening in the group that received music training. Interestingly, an enhanced P3-like late positive complex was observed in older musicians with a generator in the right auditory cortex (Zendel & Alain, 2014). This enhancement was evoked during an active listening task where participants were asked to perceptually isolate concurrently occurring sounds based on frequency cues, and the right auditory cortex is known to be sensitive to spectral features (Warrier et al., 2009; Zatorre, 1988). In the current study, the enhanced P250 was over fronto-left regions that are known to be important for understanding speech (Hickok & Poeppel, 2007). This suggests that music training improves the ability to orient attention towards salient features in the incoming auditory stimulus, by enhancing brain responses from regions responsible for processing features of the incoming acoustic stimulus. This interpretation would suggest that the electrophysiological benefit observed in the Mus group was related to an increased positivity that overlapped both P2 and P3 responses. This enhanced response likely represents enhanced stimulus classification as indexed by a P2, and enhanced ability to orient towards a stimulus as indexed by P3. Overall, this improved ability likely created a more robust neural representation of the word so that subsequent semantic processing of the word was facilitated.

## N600: A delayed N400?

The post-training enhancement to P250 observed in the group that received music training was followed by a reduction in the N600. The change in both P250 and N600 were related to each other and related to an improved ability to understand speech in noise. In a previous study from our group, using the same paradigm as the current study, we observed that the N400 was little-effected by noise in younger musicians, and was smaller in musicians compared to non-musicians (Zendel et al., 2015). This pattern of responses, paired with an enhanced P1 response in younger musicians, suggested that enhanced encoding of auditory features in noise facilitated lexical access in younger musicians (Zendel et al., 2015). Previous work on the N400 in older adults suggests that the N400 is delayed and reduced in amplitude in older adults (Kutas & Iragui, 1998). It is therefore likely that the N600 observed in the current study was a delayed N400 because the sample of participants were older. While the N600 was reduced in the Mus group from the Pre to Post training sessions, it is unlikely that this reduction was related to normal aging, as it occurred over a relatively short-time frame, and was associated with improved ability to understand speech in noise. It is therefore more likely that the reduction in N600 was due to an improvement in the processing and allocation of attentional resources towards the incoming speech sound before semantic processing or lexical access occurred. Support for this overall model is further strengthened by the fact that musicians who spent more time training also showed the greatest enhancement to the P250 and the greatest reduction to the N600.

## Summary

Music training improved the ability to understand speech-in-noise. While there were multiple benefits observed to different components of the auditory evoked response, a regression analysis suggests that the electrophysiological variables that changed with music training were highly related, and thus measuring a single factor (Earlier N100, Enhanced P2/3, Reduced N600). The pattern of results is consistent with the idea that music training improved encoding of the incoming stimulus (earlier N100); this allowed for an enhanced ability to detect salient features in the word (enhanced P250), which in turn facilitated semantic processing/lexical access of that word (reduced N600).One possible explanation for this pattern of results is based on the speech-motor system. There is a long history of work supporting the idea that speech perception relies in part on the speech-motor system (Liberman & Mattingly, 1985; Pulvermüller & Fadiga, 2010). Left frontal regions have been associated with using an articulatory model to distinguish phonemic sounds (Zatorre, Evans, Meyer, & Gjedde, 1992). Musical training may have strengthened the ability to generate an articulatory model of the incoming word, based on a more robust neural representation of the incoming word. The enhanced articulatory model of the incoming speech stimulus would then facilitate sematic processing of the incoming word. This would be reflected in an enhanced P250, and reduced N600. This model is likely as we did not observe any training-related interactions with Noise level on the event-related brain responses. This suggests that music training in older adults is not improving the ability to perceptually supress noise, but rather is related to enhanced processing of speech. One critical component of music training is connecting the auditory and motor systems, and the current results suggest that this connection may transfer to the speech perception system as well.

Overall these findings are most critical from a rehabilitation perspective. Short-term music training can have positive impact on hearing abilities. This could have a cascading effect on other cognitive abilities, as hearing difficulties typically precede other forms of cognitive decline (Lin et al., 2013). Music training is also relatively inexpensive and enjoyable, suggesting that it could be easily incorporated into many peoples lives. It remains possible that the benefits of music-training could be strengthened by identifying individual differences in susceptibility to the benefits of music-training, and by examining how different types of music-training or music-based rehabilitation contribute to enhanced auditory abilities.

## Acknowledgements

The authors would like to thank Olivier Dussault, Charles-David Tremblay, Samira Mellah and Mihaela Felezeu for assistance with data collection. Support for this research came from the Canada Research Chairs program, the GRAMMY foundation, Fondation Caroline Durant, Fonds de recherche du Québec-Santé (FRQS), and Natural Sciences and Engineering Research Council of Canada Collaborative Research and Training Experience Program in Auditory Cognitive Neuroscience (NSERC–CREATE–ACN).

## References

Alain, C., Snyder, J. S., & Dyson, B. J. (2006). Aging and the perceptual organization of Sounds: A Change of Scene? Handbook of Models for Human Aging, 759–769. https://doi.org/10.1016/B978-012369391-4/50065-5

Alain, C., Zendel, B. R., Hutka, S., & Bidelman, G. M. (2014). Turning down the noise: The benefit of musical training on the aging auditory brain. Hearing Research, 308, 162–173. https://doi.org/10.1016/j.heares.2013.06.008

Amenedo, E., & Díaz, F. (1998). Aging-related changes in processing of non-target and target stimuli during an auditory oddball task. Biological Psychology, 48 (3), 235–67. Retrieved from http://www.ncbi.nlm.nih.gov/pubmed/9788763

Anderer, P., Semlitsch, H. V, & Saletu, B. (1996). Multichannel auditory event-related brain potentials: effects of normal aging on the scalp distribution of N100, P2, N2 and P300 latencies and amplitudes. Electroencephalography and Clinical Neurophysiology, 99 (5), 458–72. Retrieved from http://www.ncbi.nlm.nih.gov/pubmed/9020805

Anderson, S., White-Schwoch, T., Parbery-Clark, A., & Kraus, N. (2013). Reversal of age-related neural timing delays with training. Proceedings of the National Academy of Sciences of the United States of America, 110 (11), 4357–62. https://doi.org/10.1073/pnas.1213555110

Billings, C. J., Tremblay, K. L., Stecker, G. C., & Tolin, W. M. (2009). Human evoked cortical activity to signal-to-noise ratio and absolute signal level. Hearing Research, 254 (1–2), 15–24. https://doi.org/10.1016/j.heares.2009.04.002

Coffey, E. B. J., Mogilever, N. B., & Zatorre, R. J. (2017). Speech-in-noise perception in musicians?: A review. Hearing Research. https://doi.org/10.1016/j.heares.2017.02.006

Crowley, K. E., & Colrain, I. M. (2004). A review of the evidence for P2 being an independent component process: age, sleep and modality. Clinical Neurophysiology?: Official Journal of the International Federation of Clinical Neurophysiology, 115 (4), 732–44. https://doi.org/10.1016/j.clinph.2003.11.021

Davis, S. W., Dennis, N. A., Daselaar, S. M., Fleck, M. S., & Cabeza, R. (2008). Que PASA? The Posterior-Anterior Shift in Aging. Cerebral Cortex, 18, 1201–1209. https://doi.org/10.1093/cercor/bhm155

Dinteren, R. Van, Arns, M., Jongsma, M. L. A., & Kessels, R. P. C. (2014). P300 Development across the Lifespan?: A Systematic Review and Meta-Analysis, 9 (2). https://doi.org/10.1371/journal.pone.0087347

Fujioka, T., Ross, B., Kakigi, R., Pantev, C., & Trainor, L. J. (2006). One year of musical training affects development of auditory cortical-evoked fields in young children. Brain: A Journal of Neurology, 129 (Pt 10), 2593–608. https://doi.org/10.1093/brain/awl247

Gates, G. A., & Mills, J. H. (2005). Presbycusis. Lancet, 366 (9491), 1111–20. https://doi.org/10.1016/S0140-6736(05)67423-5

Haxby, J. V, & Maisog, J. M. (1994). Age-related Processing Changes in Cortical Blood Flow Activation of Faces and Location during Visual, 14 (March).

Hickok, G., & Poeppel, D. (2007). The cortical organization of speech processing. Nature Reviews Neuroscience, 8 (May), 393–402.

Kaplan-Neeman, R., Kishon-Rabin, L., Henkin, Y., & Muchnik, C. (2006). Identification of syllables in noise: Electrophysiological and behavioral correlates. The Journal of the Acoustical Society of America, 120 (2), 926. https://doi.org/10.1121/1.2217567

Kok, A. (2001). On the utility of P3 amplitude as a measure of processing capacity. Psychophysiology, 38 (3), 557–577. https://doi.org/10.1016/S0167-8760(98)90168-4

Kraus, N., & Chandrasekaran, B. (2010). Music training for the development of auditory skills. Nature Reviews. Neuroscience, 11 (8), 599–605. https://doi.org/10.1038/nrn2882

Kraus, N., & White-Schwoch, T. (2015). Unraveling the Biology of Auditory Learning: A Cognitive– Sensorimotor–Reward Framework. Trends in Cognitive Sciences, xx, 1–13. https://doi.org/10.1016/j.tics.2015.08.017

Kutas, M., & Iragui, V. (1998). The N600 in a semantic categorization task across 6 decades, 108, 456–471.

Lappe, C., Herholz, S. C., Trainor, L. J., & Pantev, C. (2008). Cortical plasticity induced by short-term unimodal and multimodal musical training. The Journal of Neuroscience: The Official Journal of the Society for Neuroscience, 28 (39), 9632–9. https://doi.org/10.1523/JNEUROSCI.2254-08.2008

Liberman, A., & Mattingly, I. (1985). The motor theory of speech perception revised. Cognition, 21, 1–36.

Lin, F. R., Yaffe, K., Xia, J., Xue, Q.-L., Harris, T. B., Purchase-Helzner, E., … Simonsick, E. M. (2013). Hearing loss and cognitive decline in older adults. JAMA Internal Medicine, 173 (4), 293– https://doi.org/10.1001/jamainternmed.2013.1868

Martin, B. a, Sigal, a, Kurtzberg, D., & Stapells, D. R. (1997). The effects of decreased audibility produced by high-pass noise masking on cortical event-related potentials to speech sounds/ba/and/da. The Journal of the Acoustical Society of America, 101 (3), 1585–99. Retrieved from http://www.ncbi.nlm.nih.gov/pubmed/9069627

Mathers, C., Smith, A., & Concha, M. (2001). Global burden of hearing loss in the year 2000,(4), 1–30.

Mick, P., Kawachi, I., & Lin, F. R. (2014). The Association between Hearing Loss and Social Isolation in Older Adults. Otolaryngology--Head and Neck Surgery, 150 (3), 378–84. https://doi.org/10.1177/0194599813518021

Näätänen, R., & Picton, T. (1987). The N100 wave of the human electric and magnetic response to sound: a review and an analysis of the component structure. Psychophysiology, 24 (4), 375–425. Retrieved from http://www.ncbi.nlm.nih.gov/pubmed/3615753

Nasreddine, Z. S., Phillips, N. A., Bedirian, V., Charbonneau, S., Whitehead, V., Collin, I., … Chertkow, H. (2005). The Montreal Cognitive Assessment, MoCA: A Brief Screening. Journal of the American Geriatrics Society, 53, 695–699.

Parbery-Clark, A., Anderson, S., Hittner, E., & Kraus, N. (2012). Musical experience offsets age-related delays in neural timing. Neurobiology of Aging, 33 (7), 1483. e1-4. https://doi.org/10.1016/j.neurobiolaging.2011.12.015

Parbery-clark, A., Skoe, E., & Kraus, N. (2009). Musical experience limits the degradative effects of background noise on the neural processing of sound. The Journal of Neuroscience, 29 (45), 14100–7. https://doi.org/10.1523/JNEUROSCI.3256-09.2009

Parbery-Clark, A., Skoe, E., Lam, C., & Kraus, N. (2009). Musician enhancement for speech-in-noise. Ear and Hearing, 30 (6), 653–61. https://doi.org/10.1097/AUD.0b013e3181b412e9

Parbery-Clark, A., Strait, D. L., Anderson, S., Hittner, E., & Kraus, N. (2011). Musical experience and the aging auditory system: implications for cognitive abilities and hearing speech in noise. PloS One, 6 (5), e18082. https://doi.org/10.1371/journal.pone.0018082

Pichora-Fuller, M. K., Schneider, B. a, & Daneman, M. (1995). How young and old adults listen to and remember speech in noise. The Journal of the Acoustical Society of America, 97 (1), 593–608. Retrieved from http://www.ncbi.nlm.nih.gov/pubmed/7860836

Pulvermüller, F., & Fadiga, L. (2010). Active perception: sensorimotor circuits as a cortical basis for language. Retrieved from http://dx.doi.org/10.1038/nrn2811

Quaranta, N., Coppola, F., Casulli, M., Barulli, M. R., Panza, F., Tortelli, R., … Logroscino, G. (2015). Epidemiology of age related hearing loss: A review. Hearing, Balance and Communication, (December 2014), 1–5. https://doi.org/10.3109/21695717.2014.994869

Robert Frisina, D., & Frisina, R. D. (1997). Speech recognition in noise and presbycusis: Relations to possible neural mechanisms. Hearing Research, 106 (1–2), 95–104. https://doi.org/10.1016/S0378-5955(97)00006-3

Schneider, B. A., Pichora-Fuller, M. K., & Daneman, M. (2010). Effects of Senescent changes in audition and cognition on spoken language comprehension. In S. Gordon-Salant, R. D. Frisina, A. N. Popper, & R. R. Fay (Eds.), Aging Auditory System (pp. 167–209). New York: Springer US.

Shahin, A., Bosnyak, D. J., Trainor, L. J., & Roberts, L. E. (2003). Enhancement of neuroplastic P2 and N1c auditory evoked potentials in musicians. The Journal of Neuroscience: The Official Journal of the Society for Neuroscience, 23 (13), 5545–52. Retrieved from http://www.ncbi.nlm.nih.gov/pubmed/12843255

Shahin, A., Roberts, L. E., Pantev, C., Trainor, L. J., & Ross, B. (2005). Modulation of P2 auditory-evoked responses by the spectral complexity of musical sounds. Neuroreport, 16 (16), 1781–5. Retrieved from http://www.ncbi.nlm.nih.gov/pubmed/16237326

Stenklev, N. C., & Laukli, E. (2004). Presbyacusis—hearing thresholds and the ISO 7029. International Journal of Audiology, 43 (5), 295–306. https://doi.org/10.1080/14992020400050039

Tierney, A. T., Krizman, J., & Kraus, N. (2015). Music training alters the course of adolescent auditory development. https://doi.org/10.1073/pnas.1505114112

Warrier, C., Wong, P., Penhune, V., Zatorre, R., Parrish, T., Abrams, D., & Kraus, N. (2009). Relating structure to function: Heschl’s gyrus and acoustic processing. The Journal of Neuroscience?: The Official Journal of the Society for Neuroscience, 29 (1), 61–9. https://doi.org/10.1523/JNEUROSCI.3489-08.2009

Zatorre, R. J. (1988). Pitch perception of complex tones and human temporal-lobe function. The Journal of the Acoustical Society of America. https://doi.org/10.1121/1.396834

Zatorre, R. J., Evans, A. C., Meyer, E., & Gjedde, A. (1992). Lateralization of Phonetic and Pitch Discrimination in Speech Processing. Science, 256 (5058), 846–849.

Zendel, B. R., & Alain, C. (2012). Musicians experience less age-related decline in central auditory processing. Psychology and Aging, 27 (2), 410–7. https://doi.org/10.1037/a0024816

Zendel, B. R., & Alain, C. (2013). The influence of lifelong musicianship on neurophysiological measures of concurrent sound segregation. Journal of Cognitive Neuroscience, 25 (4), 503–16. https://doi.org/10.1162/jocn_a_00329

Zendel, B. R., & Alain, C. (2014). Enhanced attention-dependent activity in the auditory cortex of older musicians. Neurobiology of Aging, 35 (1), 55–63. https://doi.org/10.1016/j.neurobiolaging.2013.06.022

Zendel, B. R., Tremblay, C., Belleville, S., & Peretz, I. (2015). The impact of musicianship on the cortical mechanisms related to seperating speech from background noise. Journal of Cognitive Neuroscience, 27 (5).

